# Site-dependent Treg cell transcriptional reprograming in a metastatic colorectal cancer model holds prognostic significance

**DOI:** 10.1101/2025.03.13.643028

**Authors:** Sonia Aristin Revilla, Andre Verheem, Cynthia Lisanne Frederiks, Yongsoo Kim, Theofilos Chalkiadakis, Bastiaan J. Viergever, Balázs Győrffy, Enric Mocholi, Onno Kranenburg, Stefan Prekovic, Paul James Coffer

**Author notes:** Joint senior authors (Enric Mocholi, Onno Kranenburg, Stefan Prekovic, Paul James Coffer).

## Abstract

In colorectal cancer (CRC), tumor-infiltrating regulatory T (Treg) cells suppress anti-tumor immunity, promoting immune evasion and tumor progression. Effective therapies require selectively targeting tumor-infiltrating Treg (TI-Treg) cells while preserving systemic Treg cells, necessitating insight into their adaptations within the tumor microenvironment. Here, CRC-organoids were implanted in the liver of Foxp3eGFP mice to investigate location-specific phenotypic differences in TI-Treg cells. Tumor tissue exhibited an increased proportion of Treg cells and a decrease of effector CD4⁺ and CD8⁺ T cells compared to matched healthy tissue. RNA sequencing of Treg cells isolated from the spleen, primary liver tumor transplant, or metastases identified gene expression profiles previously associated with CRC-related Treg cells in patients. Location-specific differences included elevated expression of WNT-pathway genes in peritoneal TI-Treg cells compared to liver counterparts. Higher expression of genes upregulated in liver TI-Treg cells correlated with poor CRC prognosis. Splenic Treg cells from tumor-bearing mice displayed distinct transcriptional profiles from both their healthy counterparts and TI-Treg cells, suggesting they represent a distinct CD4^+^ population. Taken together, these findings highlight TI-Treg cells heterogeneity across different tumor sites and the distinct nature of splenic Treg cells in tumor-bearing hosts.

## Introduction

Colorectal cancer (CRC) is a heterogeneous disease categorized into molecular subgroups, defined by mutations and gene expression profiles, which predict clinical outcomes and therapeutic responses ^1^. CRC tumors shape the tumor microenvironment (TME) to establish a immunosuppressive niche that controls immune cell infiltration, enabling immune escape and tumor progression ^2–4^. Understanding the types, frequencies, and distribution of immune cells in the TME is crucial for gaining insight into tumor-immune interactions and improving patient diagnosis and prognosis ^5,6^.

Tumor-infiltrating lymphocytes (TILs), enriched among the tumor-infiltrating immune cells in CRC, are crucial for molecular classification, prognosis, and immunotherapy response ^2,7,8^. The Immunoscore system classifies CRC tumors as “hot,” “altered,” or “cold” based on TILs, providing independent prognostic value and linking molecular classification to the immune environment ^2,6,7,9^. High microsatellite instability CRC tumors, characterized by defective DNA mismatch repair, have a high mutational burden and neoantigen production. This often results in significant immune infiltration, including cytotoxic CD8^+^ T cells and CD4^+^ Th1 cells, leading to a “hot” classification and better immunotherapy responses ^9–11^. In contrast, most CRC tumors have proficient DNA mismatch repair and microsatellite stability or low instability have low mutation rates, minimal T cell infiltration, and poor immunotherapy response, resulting in a “cold” classification and worse prognosis ^9,10^. “Altered” CRC tumors , including “excluded” and “immunosuppressed” types, exhibit intermediate immune infiltration and poor immunotherapy responses ^9^. “Excluded” tumors have T cells restricted to the margins, while “immunosuppressed” tumors feature minimal T cell infiltration, with T regulatory (Treg) cells predominating ^9^.

Treg cells are an immunosuppressive subset of CD4^+^ T cells crucial for maintaining immunological homeostasis and self-tolerance ^12,13^. They are characterized by expression of the transcription factor FOXP3 (forkhead box P3), which regulates their development and function in both mice and humans ^12,13^. Treg cells exhibit phenotypic plasticity, allowing them to co-express various transcription factors, surface markers, and cytokines, and adopt different phenotypes in response to environmental changes ^4,8,14,15^. In human disease, two CRC FOXP3^+^ T cell subpopulations have been identified: suppressive FOXP3^high^ Treg cells, linked to poorer outcomes and lower disease-free survival, and non-suppressive FOXP3^low^ T cells, associated with better prognosis and a pro-inflammatory environment ^16^. FOXP3 expression can occur in activated effector T cells, which are not functionally suppressive ^17^.

In CRC, Treg cells constitute a high proportion of CD4^+^ T cells in the TME compared to healthy colon tissue, with late-stage patients showing a higher Treg-to-effector T (Teff) cell ratio, indicative of a more immunosuppressive environment ^8,18,19^. This increased accumulation of suppressive TI-Treg cells is linked to tumor progression, metastasis, and immunotherapy failure, contributing to a poorer prognosis ^4,20–24^. The plasticity and heterogeneity of tumor-infiltrating Treg (TI-Treg) cells offer diverse mechanisms to inhibit effective anti-tumor immunity, thereby promoting immune evasion ^4,8,15^. Therefore, enhancing current immunotherapy strategies by targeting TI-Treg cells holds significant potential. Approaches such as low-dose cyclophosphamide treatment or monoclonal antibodies targeting checkpoint receptors have shown promise in depleting TI-Treg cells and activating anti-tumor immune responses in CRC patients ^4,25,26^. However, the specific characteristics and functional mechanisms of these cells remain unclear, including their phenotypic markers, tissue residency, and the cytokine environment that influences their activity. This lack of clarity complicates efforts to selectively target TI-Treg subsets while preserving beneficial immune responses. Therefore, effectively distinguishing and targeting these subsets is essential for improving treatment outcomes in CRC.

Exploring the identity and heterogeneity of CRC TI-Treg cells is crucial, as it significantly impacts patient outcomes and the effectiveness of targeted therapies. Understanding the diverse phenotypes and functional states of TI-Treg cells can enhance prognostic accuracy and inform the development of more effective treatment strategies. Here, we show that TI-Treg cells exhibit distinct phenotypes at different locations and assessed the systemic impact of tumor development on Treg cell transcriptional profiles. To this end, we used a novel *in vivo* tumor-organoid implantation model to evaluate the transcriptional profile of splenic and tumor-infiltrating Treg cells. This model provides insights into Treg cell diversity at different tumor sites and increases our understanding of their prognostic significance, which is critical for identifying viable targets for immunotherapy and developing tailored approaches for the selective depletion of TI-Treg cells.

## Results

### Increased numbers of TI-Treg cells at CRC metastatic sites

To characterize tumor-infiltrating immune cells at CRC metastatic sites, we used an immunocompetent Foxp3EGFP mouse model. Here, CRC tumor-organoids, expressing firefly luciferase, were embedded in collagen droplets and micro surgically implanted into the liver, mimicking the most common site of CRC metastasis. As we have previously shown, liver implantation results in the development of CRC metastatic tumors at distinct sites ^27^. Using non-invasive bioluminescence imaging (BLI), we monitored tumor development and metastasis. Animals were euthanized at humane endpoint due to tumor growth or signs of discomfort. Metastatic tumors were identified postmortem through visual inspection and BLI measurements (**Figure 1a**). Within two weeks, 76% of mice developed tumors at the implantation sites in the liver, termed “primary liver tumor transplant”, with metastatic tumors emerging thereafter in 92% of these mice (see **Table 1**). The spleen, and visually detected tumors from tumor-bearing mice, were isolated, and some were retained for histological tissue analysis. Additionally, the spleen from healthy mice, as well as healthy tissue from the corresponding organs where tumors developed in tumor-bearing mice, were isolated and used as controls. We observed a similar size and weight of the spleens from tumor-bearing mice compared to healthy spleens (**Suppl. Figure 1a-b**). To assess immune cell infiltration in both tumor and healthy tissue, we performed immunohistochemistry (IHC) staining for CD45-positive cells. A significant decrease of CD45^+^ immune cells per mm^2^ was observes in the spleen (15196 ± 1100) and liver tumors (1980 ± 666) of tumor-bearing mice compared to healthy spleen (17544 ± 954) and liver (2585 ± 1812) controls (**Figure 1b**). In contrast, CD45^+^ immune cells per mm^2^ were significantly increased in metastatic lung (10418 ± 4679) and peritoneal (3231 ± 1031) tumors relative to healthy lung (3602 ± 1661) and peritoneum (497 ± 378) controls. Immune cell localization displayed distinct pattern, with immune cells clustering at the tumor margins and less dense regions in liver and peritoneal tumors, whereas in lung tumors, immune cells are more uniformly distributed throughout the tumor tissue (**Suppl. Figure 1c**).

**Figure 1.**
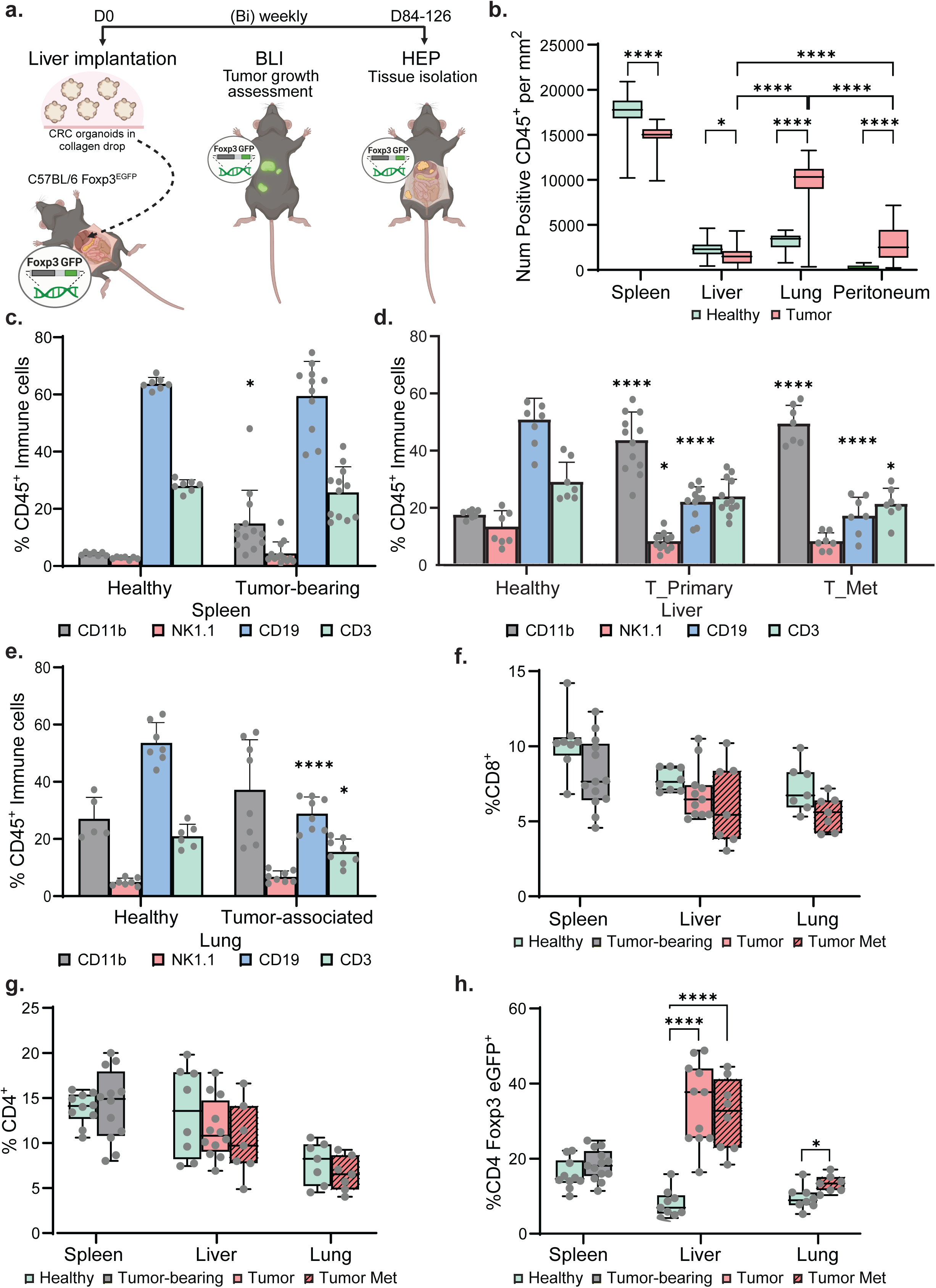
Increased immunosuppressive tumor-infiltrating immune cells at CRC metastatic sites. (**a**) Schematic timeline illustrating liver (primary) implantation, tumor growth monitoring by bioluminescence imaging (BLI), and postmortem tumors identification. (**b**) Quantification of CD45^+^ cells per mm2 in the spleen, liver (healthy versus tumors), lung (healthy versus tumors), and peritoneum (healthy versus tumors) of healthy versus tumor-bearing mice. (**c-e**) Percentage of myeloid cells (CD45^+^ CD11b^+^ cells), NK cells (CD45^+^ NK1.1^+^ cells), B cells (CD45^+^ CD19^+^ cells) and T cells (CD45^+^ CD3^+^ cells) in the (**c**) spleen, (**d**) liver (healthy versus primary or metastatic tumors), and (**e**) lung (healthy versus tumor-associated) of healthy versus tumor-bearing mice, assessed by flow cytometry.(**f-h**) Percentage of (**f**) CD8^+^ T cells, (**g**) CD4^+^ T cells, and (**h**) CD4^+^ Foxp3 eGFP^+^ Treg cells in the spleen, liver (healthy compared to primary or metastatic tumors), and the lung (healthy compared to tumor-associated) of healthy versus tumor-bearing mice, assessed by flow cytometry. Data are represented as mean ± SD. P-values were calculated using one-way ANOVA and unpaired t-test analysis. *p ≤ 0.05, **p ≤ 0.01, ***p ≤ 0.001 and ****p ≤ 0.0001.

**Table1.** Number of mice developing primary tumors and metastasis.

| **Mice** | **Implanted** | **Primary liver tumor transplant** | **Liver met** | **Peritoneal met** | **Lung met** | **Other intra-abdominal met** |
| --- | --- | --- | --- | --- | --- | --- |
| **Number** | 17 | 13 | 12 | 11 | 4 | 4-9 |
| **Developed tumor (%)** |  | 76% | 71% | 65% | 24% | 35-53% |
| **Developed metastasis (%)** |  |  | 92% | 85% | 31% | 46-69% |

Tumors and healthy tissue samples were subsequently processed into single-cell suspensions to further assess the abundance of different CD45^+^ immune cell populations by flow cytometry. Immune profiling was evaluated by flow cytometry analyzing the presence of myeloid cells (CD45^+^ CD11b^+^ cells), NK cells (CD45^+^ NK1.1^+^ cells), B cells (CD45^+^ CD19^+^ cells) and T cells (CD45^+^ CD3^+^ cells) (**Suppl. Figure 1d**). A significant increase in the percentage of myeloid cells was observed in the spleens (14.9% ± 11.1%) and liver tumors, both primary transplant (42.9% ± 9.5%) and metastatic (48.7% ± 6%), of tumor-bearing mice, compared to healthy spleen (4.2% ± 0.5%) and liver (16.9% ± 1.3%) tissues (**Figure 1c-d**). Similarly, an elevated percentage of myeloid cells was detected in lung metastatic sites (37.2% ± 16.3%) compared to healthy controls (27.1% ± 6.7%) (**Figure 1e**). In contrast, the relative percentages of NK cells, B cells, and T cells were generally reduced at the different tumor sites compared to their paired controls (**Figure 1c-e**), with no significant difference observed in the spleen of tumor-bearing mice compared to healthy spleen (**Figure 1c**). To further characterize the tumor-infiltrating lymphocyte (TIL) populations, CD8^+^ T cells, CD4^+^ T cells, and CD4^+^ Foxp3 eGFP^+^ Treg cells within the CD45^+^ CD3^+^ T cell population were analyzed (**Suppl. Figure 1e**). Both CD8^+^ and CD4^+^ T cell populations showed a consistent decrease in percentage across the different tumor sites compared to their paired controls (**Figure 1f-g**). In contrast, the percentage of CD4^+^ Foxp3 eGFP^+^ Treg cells significantly increased at all tumor sites —liver primary tumor transplant (34.6% ± 10.4%), liver metastasis (31.7% ± 9%), and lung metastasis (13.3% ± 2.2%)— compared to healthy liver (8.1% ± 3.5%) and lung (9.5% ± 3%) tissues. Notably, TI-Treg cell levels were nearly doubled in liver tumors compared to lung metastases. Treg cell percentages in the spleen did not significantly differ between tumor-bearing and healthy mice (**Figure 1h**). However, TI-Treg cell levels were nearly doubled in liver tumors (34.6% ± 10.4%) compared to lung (13.3% ± 2.2%) or peritoneal (22.6% ± 5%) metastases (**Suppl. Figure 1f**). Taken together, these observations demonstrate an increased presence of myeloid cells and TI-Treg cells while a decrease in CD4^+^ and CD8^+^ T cells in metastatic sites, which may be associated with an immunosuppressive TME in CRC metastases.

### Treg cell transcriptional profiles vary by location

To define the transcriptional identity of TI-Treg cells at the different locations, we performed bulk RNA-sequencing (**Figure 2a**). We utilized a staged cell sorting approach to isolate CD4^+^ Foxp3 eGFP^neg^ T cells and CD4^+^ Foxp3 eGFP^+^ splenic Treg cells from the spleens of healthy mice. Additionally, CD4^+^ Foxp3 eGFP^+^ splenic Treg cells of tumor-bearing animals were isolated, as well as CD4^+^ Foxp3 eGFP^+^ TI-Treg cells from tumor sites including the primary liver tumor transplant, liver metastases, and peritoneal metastases (**Figure 2a**). As anticipated, Principle Component Analysis (PCA) showed that splenic eGFP^neg^ CD4^+^ T cells and splenic CD4^+^ Foxp3 eGFP^+^ Treg cells from healthy and tumor-bearing mice were distinct populations from CD4^+^ Foxp3 eGFP^+^ TI-Treg cells in the CRC metastases. Additionally, Treg cells from the spleen of tumor-bearing mice clustered separately from those in healthy mice (**Figure 2b**). To assess sample similarity based on transcriptional profiles, unsupervised clustering was performed, which grouped samples according to their tissue origin (**Figure 2c, Suppl. Figure 2a**). To understand these differences, Gene Ontology (GO) term analysis was conducted to delineate the biological processes involved. This analysis revealed downregulation of gene expression pathways including protein translation, and upregulation of metabolic pathways, such as oxidative phosphorylation, as well as pathways associated with cytokine production, chemotaxis and SMAD-mediated gene regulation in the TI-Treg cells compared to the different splenic CD4^+^ T cell populations (**Suppl. Figure 2b**). To evaluate the expression of Treg cell signature genes across the different CD4^+^ T cell populations, we compiled a core Treg cell signature gene list based on previously published studies ^4,28,29^. All Treg cells groups exhibited an increased expression of core Treg cell signature genes when compared to the control group of eGFP^neg^ CD4^+^ T cells (**Figure 2d, Suppl. Figure 2c**). Moreover, TI-Treg cells had elevated expression of these genes when compared to splenic Treg cells from both healthy and tumor-bearing mice (**Figure 2d, Suppl. Figure 2c**). Additionally, analysis of Treg-associated transcription factors (TFs) indicated higher expression in TI-Treg cells compared to splenic Treg cells from both healthy and tumor-bearing mice (**Figure 2e, Suppl. Figure 2d**) ^30,31^. To understand how these transcriptional differences may affect local Treg cell function, we compared Treg-associated pathways across all CD4^+^ T cell groups. TI-Treg cells upregulate genes involved in immune response regulation (TGFβ, IL-2, IL-10, CTLA4, and FOXP3 target genes), chemokine activity, and cellular metabolism (oxidative phosphorylation and the pentose phosphate pathway) (**Figure 2f**) compared to the other CD4^+^ T cell groups. Finally, to investigate FOXP3-mediated gene regulation, we analyzed the expression of a panel of defined FOXP3 target genes and found increased levels in TI-Treg cells compared to the different splenic CD4^+^ T cell populations (**Figure 2g, Suppl. Figure 2e**). Overall, these observations demonstrate that the evaluated Treg cell groups are transcriptionally distinct from each other.

**Figure 2.**
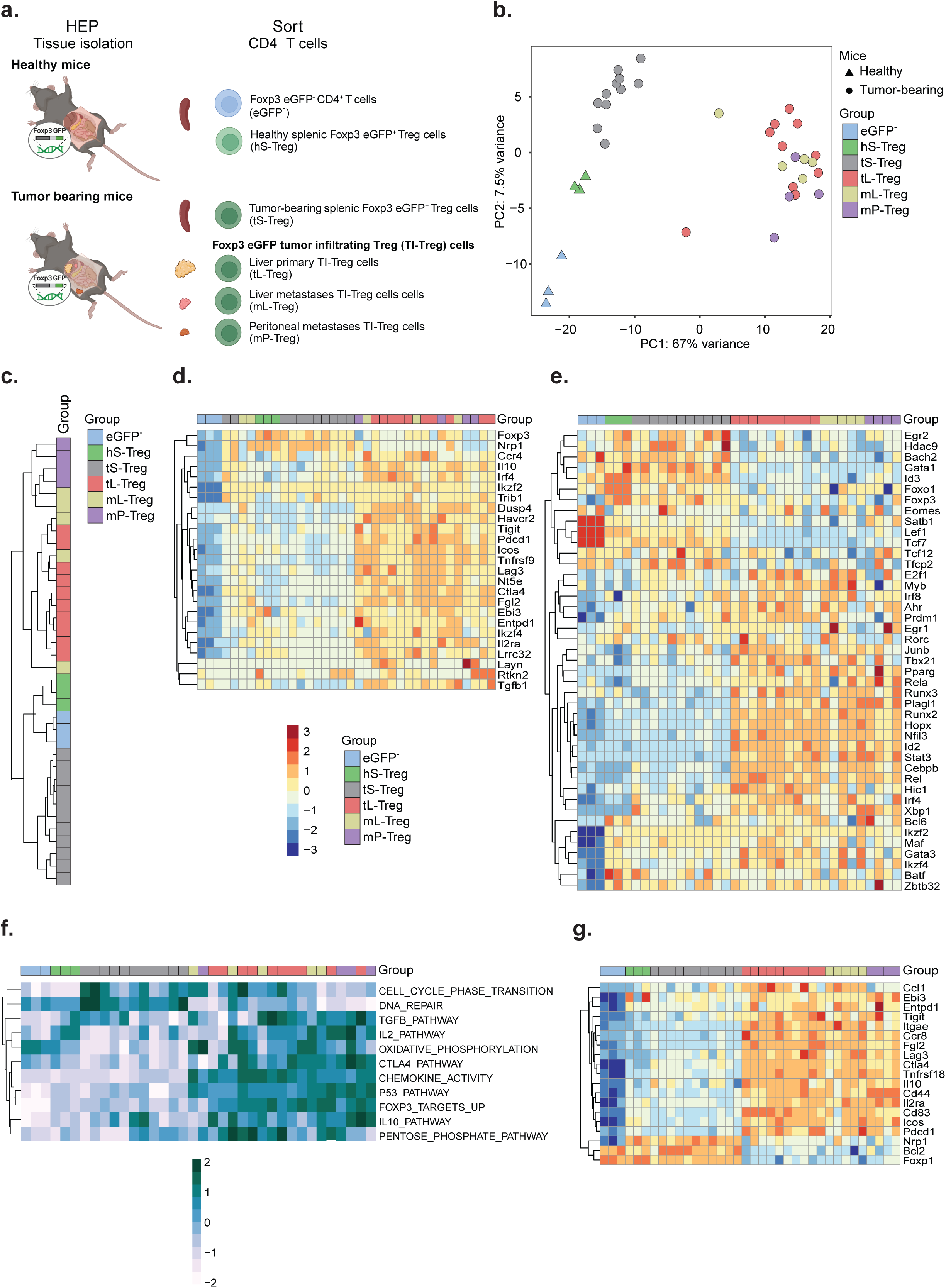
Transcriptionally distinct Treg cell subsets across different tissue location. (**a**) Schematic of isolation of different CD4^+^ T cell subsets via cell sorting from various tissues in healthy and tumor-bearing mice for bulk RNA sequencing. (**b**) PCA of healthy splenic CD4^+^ eGFP^−^ T cells, healthy splenic Treg cells, tumor-bearing splenic Treg cells, and TI-Treg cells from primary liver tumor transplants, liver metastases, and peritoneal metastases. (**c**) Unsupervised cluster analysis of all samples. (**d**) Heatmap of core Treg cells signature genes expression across all groups. (**e**) Heatmap of Treg-associated TF gene expression across all groups. (**f**) Pathway analysis focus on Treg-associated pathways across all groups. (**g**) Heatmap of Foxp3 target gene expression across all groups.

### Tumor location defines TI-Treg cell transcriptome

To compare transcriptional similarities between eGFP^+^ TI-Treg cells with CRC TI-Treg cells in patients, we analyzed genes reported to be upregulated in CRC TI-Treg cells from patients relative to peripheral Treg cells in healthy tissues or peripheral blood ^4,32^. We observed significant upregulation of these genes in all eGFP^+^ TI-Treg cells compared to splenic Treg cells (**Figure 3a-b**). eGFP^+^ TI-Treg cells showed increased expression of markers associated with activation and immune suppression, including *Lag3* (LAG-3), *Havcr2* (TIM-3), *Icos* (ICOS), *Ctla4* (CTLA-4), *Pdcd1* (PD-1), *Pdcd1lg2* (PD-L2), *Tnfrsf4* (OX40), *Tnfrsf18* (GITR), *Tnfrsf9* (4-1BB), and *Tigit* (TIGIT) ^28,29^. Additionally, we also observed a significant upregulation of key immunoregulatory genes, including *Tgfb1* (TGF-β), *Gzmb* (Granzyme B), *Il10* (IL-10), and *Vegfa* (VEGF) ^28,29^. To further characterize the transcriptional signatures of eGFP^+^ TI-Treg cells, we conducted a differential gene expression analysis comparing eGFP^+^ TI-Treg cells from the primary liver tumor transplants with those from metastatic locations. Minimal differences were observed between eGFP^+^ TI-Treg cells from the primary liver tumor transplants and liver metastases as might be expected (**Figure 3c**). However, a comparison between eGFP^+^ TI-Treg cells from the peritoneal metastases compared to those in primary liver tumor transplants identified 269 upregulated and 187 downregulated differentially expressed genes (DEGs) (p ≤ 0.05, log-fold change ≥ 2) (**Figure 3d**, **Suppl. Table 1**). To define biological processes associated with these DEGs, we performed pathway analysis, which identified their involvement in Wnt signaling (**Figure 3e**). Gene Set Enrichment Analysis of these DEGs further confirmed significant enrichment of Wnt signaling pathway in eGFP^+^ TI-Treg cells from peritoneal metastases compared to those from primary liver tumor transplants (**Figure 3f**). In summary, differential gene expression analysis revealed distinct transcriptional signatures of eGFP^+^ TI-Treg cells between liver and peritoneal tumors, highlighting key differences in Wnt signaling pathway.

**Figure 3.**
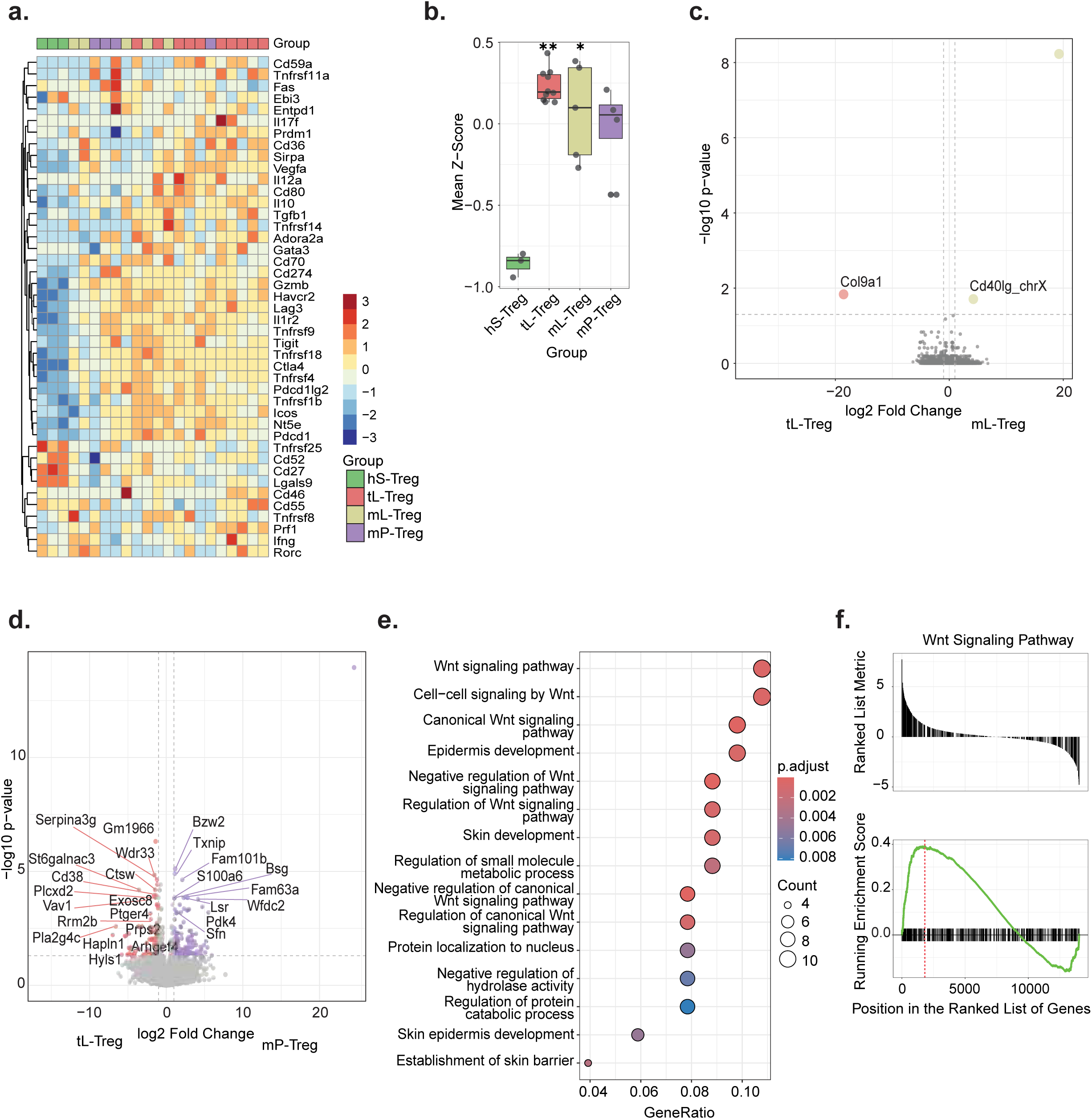
Transcriptionally distinct TI-Treg cell subsets in peritoneum and liver tumors. (**a**) Heatmap and (**b**) box plot (mean ± SD) of signature z-score for genes associated with *in vivo* CRC TI-Treg from patients cells, comparing expression in healthy splenic Treg cells and TI-Treg cells from primary liver tumor transplants, liver metastases, and peritoneal metastases. (**c-d**) Volcano plots showing log2 fold change against statistical significance for gene expression comparing TI-Treg cells from primary liver tumor transplants to (**c**) liver metastases and (**d**) peritoneal metastases (ANOVA, FDR, p ≤ 0.05). (**e**) GSEA of the DEGs between TI-Treg cells from primary liver tumor transplants and peritoneal metastases. (**f**) Pathway analysis of biological processes associated with upregulated genes in TI-Treg cells from peritoneal metastases compared to TI-Treg cells from primary liver tumor transplants. Data are represented as mean ± SD. Boxplots were analyzed with non-parametric test (two-tailed). P-values were calculated using one-way ANOVA and unpaired t-test analysis. *p ≤ 0.05, **p ≤ 0.01, ***p ≤ 0.001 and ****p ≤ 0.0001.

### Gene expression signature of TI-Treg cells has prognostic relevance

To evaluate differences in transcriptional profiles between eGFP^+^ Treg cells from the primary liver tumor transplant, and eGFP^+^ splenic Treg cells, we conducted a differential gene expression analysis. A total of 5524 genes were identified as significantly differentially expressed between TI-Treg cells from primary liver tumor transplants and splenic Treg cells (p ≤ 0.05, with 1803 upregulated and 366 downregulated genes with log-fold change ≥ 2) (**Figure 4a**, **Suppl. Table 2**). Gene Ontology (GO) term pathway analysis of upregulated genes in TI-Treg cells from primary liver tumor transplants, highlighted significant involvement of pathways regulating immune responses (**Figure 4b**). These include immune effector processes, cell activation, cytokine production, and leukocyte proliferation, as well as chemotaxis, cell migration, and adhesion. In contrast, the downregulated genes in TI-Treg cells from primary liver tumor transplants were linked to T cell activation and differentiation, as well as to processes involved in gene regulation, including translation, ribosome biogenesis, and chromatin remodelling (**Figure 4c**). To evaluate the prognostic potential of TI-Treg cell gene signatures from primary liver tumor transplants (adjusted p-value < 0.05; log-fold change ≥ 1) for CRC patient outcomes, we identified 2152 significantly upregulated genes and cross-referenced them with the single-cell CRC cohort atlas ^33^, comparing them to tumor cells. This led to the identification of 295 genes (**Suppl. Table 3**) that are expressed only in CRC lymphocytes but not in tumor epithelial cells, hereafter referred to as the CRC-T-reg signature (adjusted p-value < 0.05; log-fold change ≥ 1). We proceeded by using the CRC-T-reg signature in a large cohort (KMplotter composite cohort ^34^) of CRC containing transcriptome data and patients’ overall survival status (1061 patients). The Kaplan–Meier analysis demonstrated that the CRC-T-reg signature was able to stratify patients into good and poor prognosis groups (**Figure 4d**, p-value = 0.0025), particularly demonstrating that high expression of these genes was significantly associated with reduced overall survival in CRC. Taken together, TI-Treg exhibit a specific transcriptional program that has potential prognostic value in CRC.

**Figure 4.**
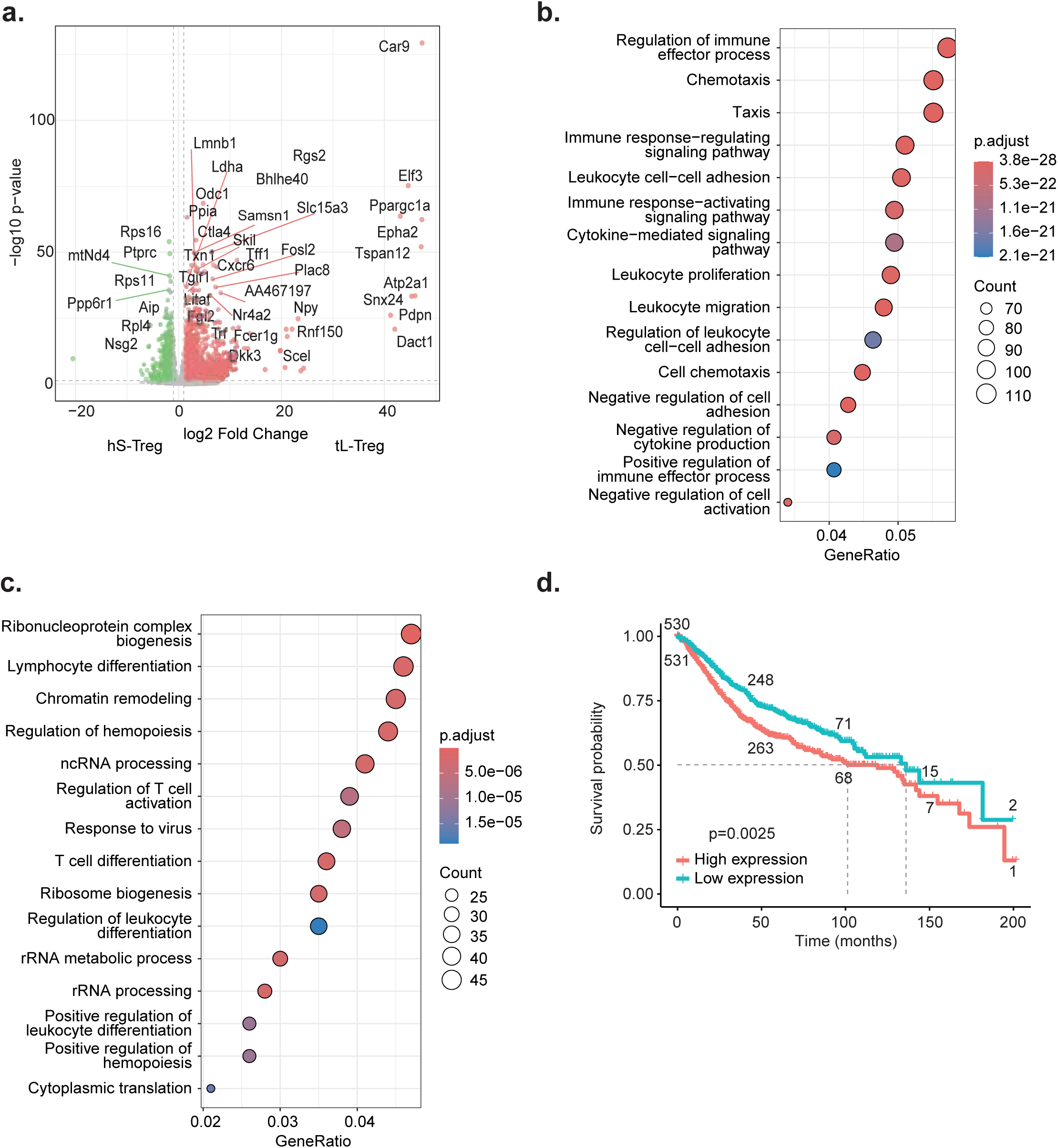
Liver TI-Treg cell gene expression patterns have prognostic relevance. (**a**) Volcano plots showing log2 fold change against statistical significance for gene expression comparing TI-Treg cells from primary liver tumor transplants to healthy splenic Treg cells (ANOVA, FDR, p ≤ 0.05). (**b-c**) Pathway analysis of biological processes associated with (**b**) upregulated and (**c**) downregulated genes in TI-Treg cells from primary liver tumor transplants compared to healthy splenic Treg cells. (**d**) Kaplan-Meier curve depicting overall survival in CRC patients stratified into low- and high-expression group based on mean expression of the CRC-T-reg signature identified from primary liver tumor transplants. The numbers on the curves represent the number of CRC patients at each time point. Data are represented as mean ± SD. P-values were calculated using unpaired t-test analysis. *p ≤ 0.05, **p ≤ 0.01, ***p ≤ 0.001 and ****p ≤ 0.0001.

### Tumor-bearing splenic Treg cells are a distinct CD4^+^ population

To assess whether in tumor-bearing mice, Treg cells exhibit a systemic tumor-associated phenotype, we compared the transcriptional profiles of splenic Treg cells from healthy and tumor-bearing animals. PCA illustrated that tumor-bearing splenic Treg cells form a distinct cluster, separate from healthy splenic Treg cells (**Figure 5a**). To determine differences between healthy and tumor-bearing splenic Treg cells, we conducted a differential gene expression analysis. This analysis identified 3569 significantly DEGs between tumor-bearing and healthy splenic Treg cells with 1078 upregulated and 75 downregulated genes with log-fold change ≥ 2 (**Figure 5b, Suppl. Table 4**). Pathway analysis of the upregulated genes in tumor-bearing splenic Treg cells highlighted their significant roles in immune responses pathways, including the regulation of innate immune responses, activation of signaling pathway and cell surface receptors, and lymphocyte proliferation (**Figure 5c**). We evaluated the expression of genes reported to be upregulated in CRC TI-Treg cells from patients, relative to peripheral Treg cells from healthy tissues or peripheral blood, in both healthy and tumor-bearing splenic eGFP^+^ Treg cells ^4,32^. In general, we observed an increase in the expression of these genes in tumor-bearing splenic Treg cells compared to healthy splenic Treg cells (see **Figure 3a-b**; **Figure 5d-e**). Additionally, we assessed the expression of the CRC-Treg signature genes (*288 that were found in the KMplotter cohort*) within healthy and tumor-bearing splenic Treg cells, finding a higher expression of this signature in tumor-bearing splenic Treg cells (see **Figure 4d**, **Figure 5f-g**). These findings demonstrate significant molecular changes in splenic Treg cells from tumor-bearing animals, suggesting a systemic response to tumor development.

**Figure 5.**
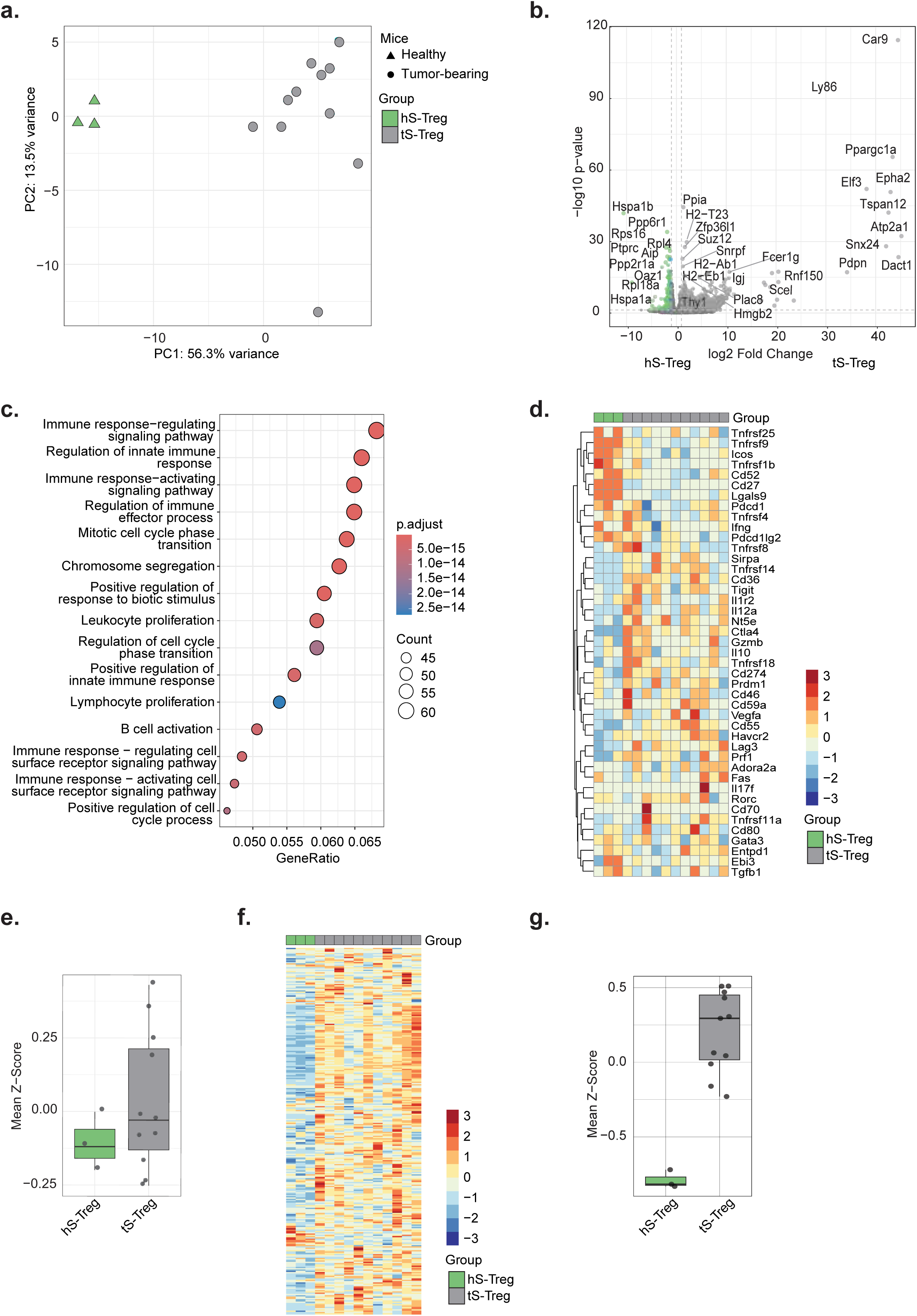
Tumor-bearing splenic Treg cells are transcriptionally more similar to CRC TI-Treg cells than their healthy counterparts. (**a**) PCA of healthy splenic Treg cells and tumor-bearing splenic Treg cells. (**b**) Volcano plots showing log2 fold change against statistical significance for gene expression comparing healthy splenic Treg cells and tumor-bearing splenic Treg cells (ANOVA, FDR, p ≤ 0.05). (**c**) Pathway analysis of biological processes associated with upregulated genes in tumor-bearing splenic Treg cells compared to healthy splenic Treg cells. (**d**) Heatmap and (**e**) box plot (mean ± SD) of signature z-score for genes associated with *in vivo* CRC TI-Treg from patients cells, comparing expression in healthy splenic Treg cells and tumor-bearing splenic Treg cells. (**f**) Heatmap and (**g**) box plot (mean ± SD) displaying the TI-Treg cell gene signature identified from primary liver tumor transplants with prognostic significance, comparing tumor-bearing splenic Treg cells compared to healthy splenic Treg cells. Data are represented as mean ± SD. Boxplots were analyzed with non-parametric test (two-tailed). P-values were calculated using unpaired t-test analysis. *p ≤ 0.05, **p ≤ 0.01, ***p ≤ 0.001 and ****p ≤ 0.0001.

To evaluate whether tumor-bearing splenic Treg cells represent an independent population or a mixture of healthy splenic Treg cells and TI-Treg cells, we conducted a comparative transcriptional analysis. First, we performed a PCA analysis, in which we generated and projected 1000 random mixtures of healthy splenic Treg cells and TI-Treg cells. These “random” samples positioned between healthy splenic and TI-Treg cells but did not cluster near the tumor-bearing splenic Treg cells (**Figure 6a**). We subsequently performed deconvolution analysis to quantify the extent of tumor-bearing splenic Treg cell profiles that could be explained by a combination of healthy splenic Treg cell and TI-Treg cell profiles. Taking profiles of healthy splenic Treg cells and TI-Treg cells as a signature, non-negative least square (NNLS) deconvolution yielded a near-perfect reconstruction of the pseudo-mixtures (close-to-one coefficient of determination; **Figure 6b**). In contrast, the performance for tumor-bearing splenic Treg cells was approximately 0.85, indicating the uniqueness of tumor-bearing splenic Treg gene expression profiles (**Figure 6b**). Taken together, these data suggest that tumor-bearing splenic Treg cells are a unique population with a distinct transcriptional profile.

**Figure 6.**
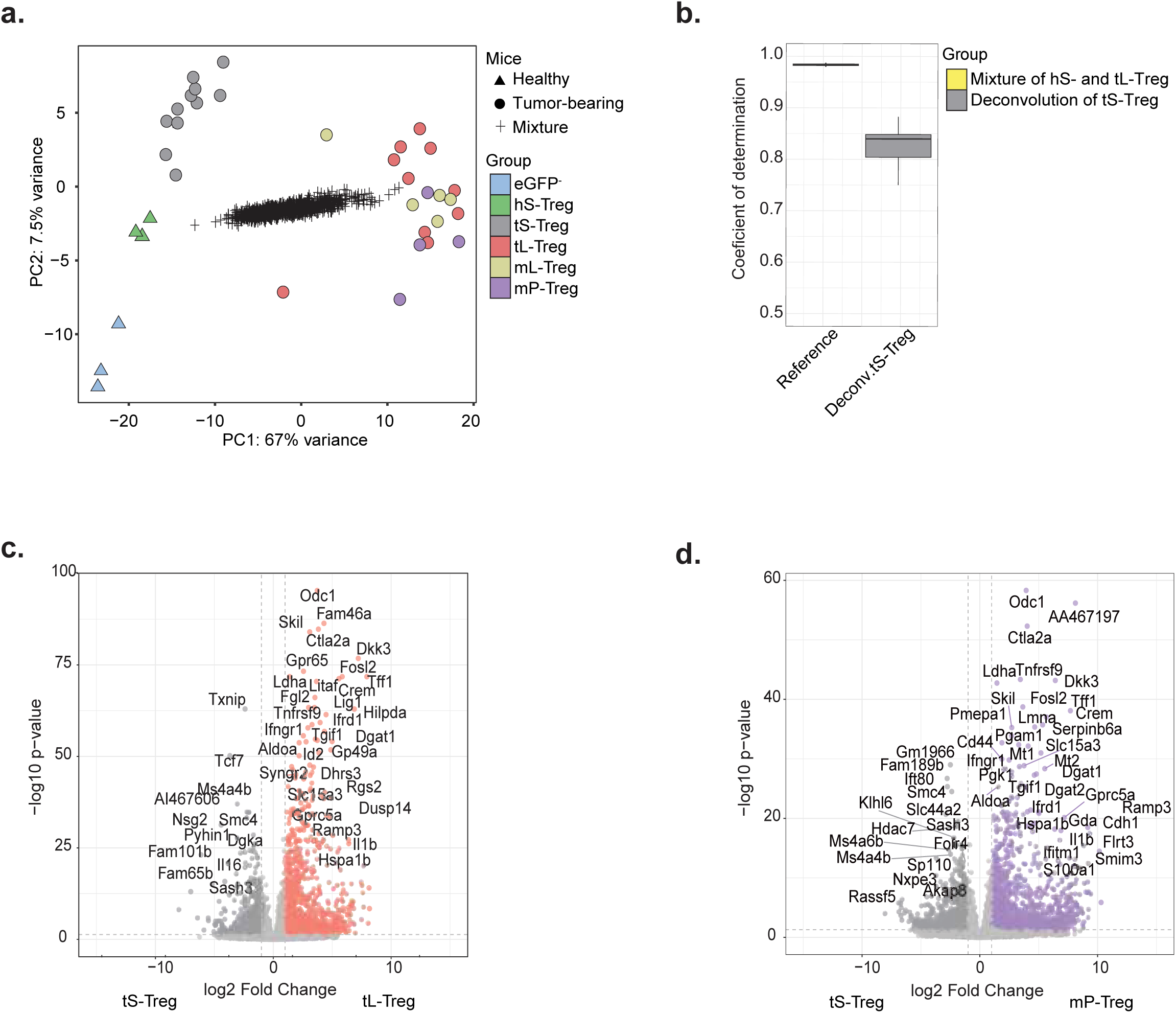
Tumor-bearing splenic Treg cells are a distinct population. (**a**) PCA of healthy splenic CD4^+^ eGFP^−^ T cells, healthy splenic Treg cells, tumor-bearing splenic Treg cells, and TI-Treg cells from primary liver tumor transplants, liver metastases, peritoneal metastases, and 1000 random mixtures of healthy splenic Treg cells and TI-Treg cells in primary liver tumor transplants. (**b**) Predicted and measured regression performance (coefficient of determination) in assessing tumor-bearing splenic Treg cells similarities to healthy splenic Treg cells and TI-Treg cells in primary liver tumor transplants. (**c**) Volcano plots showing log2 fold change against statistical significance for gene expression comparing tumor-bearing splenic Treg cells and TI-Treg cells in primary liver tumor transplants (ANOVA, FDR, p ≤ 0.05). (**d**) Volcano plots showing log2 fold change against statistical significance for gene expression comparing tumor-bearing splenic Treg cells and TI-Treg cells in peritoneal metastases (ANOVA, FDR, p ≤ 0.05).

Finally, we conducted a differential gene expression analysis to identify genes specially associated with TI-Treg cells by comparing TI-Treg cells from primary liver tumor transplants or peritoneal metastasis with tumor-bearing splenic Treg cells. Differential expression of 2505 significantly upregulated genes in TI-Treg cells from primary liver tumor transplants (p ≤ 0.05, 1449 genes with log-fold change ≥ 2), and 2458 significantly upregulated genes in TI-Treg cells from peritoneal metastasis were identified relative to tumor-bearing splenic Treg cells (p ≤ 0.05, 965 genes with log-fold change ≥ 2) (**Figure 6c-d, Suppl. Table 5 - 6**). Overall, these results highlight the value of considering tumor-bearing splenic Treg cells as a reference for identifying markers that specifically target TI-Treg cells.

## Discussion

In the CRC TME, the accumulation of suppressive TI-Treg cells is linked to poor prognosis, as these cells hinder immune surveillance and contribute to tumor progression and immunotherapy failure ^4,20–24^. While Treg cells are enriched in CRC tumors and exhibit potent immunosuppressive activity, there is still much to understand concerning their precise phenotype and mechanisms of action. Therefore, characterization of CRC TI-Treg cell identity and heterogeneity is essential for refining prognostic assessments and developing targeted therapeutic strategies. In this study, we isolated Treg cells using eGFP^+^ FOXP3 expression. In humans FOXP3 is also found in non-suppressive activated T effector cells, complicating Treg cell characterization in CRC. In mice, Foxp3 is exclusively expressed in Treg cells. This makes the Foxp3EGFP mouse model particularly valuable for specifically isolating Treg cells. To model CRC metastasis, CRC tumor organoids were implanted into the liver of immunocompetent Foxp3EGFP mice, allowing us to study immune infiltration in metastatic tumors. TI-Treg cells were specifically enriched at metastatic sites, with distinct transcriptional profiles compared to splenic Treg cells and variations based on tumor location, such as liver versus peritoneal tumors. High expression of certain TI-Treg cell genes in liver tumors correlated with shorter overall survival (**Figure 4f**), emphasizing their prognostic relevance. Moreover, tumor-bearing splenic Treg cells were also identified as a distinct population.

CRC tumors manipulate the surrounding microenvironment to induce inflammation and promote immune cell infiltration, actively shaping the TME to facilitate immune evasion ^3,35^. The immune composition within CRC tumors is highly heterogeneous and responsive to microenvironment changes ^3,35–37^. In our study, CD45^+^ immune cells were significantly reduced in spleen and liver tumors but increased in lung and peritoneal metastases in tumor-bearing mice compared to controls. Flow cytometry analysis revealed an increased proportion of myeloid cells and a decline in NK cells, B cells, and T cells, with an increase in Treg cells and a reduction in CD4^+^ and CD8^+^ T cells in tumor-bearing mice compared to controls. TI-Treg cells were more abundant in primary liver tumor transplants than in lung or peritoneal metastases, consistent with previous findings of Treg enrichment in primary tumors ^38^. These data suggest that all tumor sites were dominated by immune-suppressive cells, such as myeloid cells and Treg cells, likely promoting immune evasion. Liver tumors, however, exhibited a more immunosuppressive TME with lower immune infiltration and a higher proportion of TI-Treg cells compared to lung or peritoneal metastases (**Figure 1b, Suppl. Figure 1e**). These findings align with reports of decreased CD45^+^ immune cells, increased myeloid and TI-Treg cells, and reduced NK cells in CRC liver metastases compared to adjacent healthy liver ^36^. High TI-Treg cell levels correlate with lower overall and relapse-free survival in CRC liver metastases, while high ratios of TI-Treg to CD4^+^ and CD8^+^ T cells predict shorter survival after tumor resection ^24,39^.

Treg cells exhibit phenotypic plasticity, co-expressing various transcription factors, markers, and cytokines. This adaptability to environmental cues contributes to their heterogeneity and diverse immunosuppressive functions ^4,8,15,37,40^. RNA-seq analysis reveals distinct transcriptional phenotypes in Treg cells from distinct locations, particularly evident between spleen and tumor, as highlighted by PC1 in the PCA pot (**Figure 2b**). This distinction reflects site-specific transcriptional profiles, reflecting their adaptive roles in different microenvironments. TI-Treg cells upregulated genes related to immune suppression (e.g., *Il10*, *Tgfb1, Ctla4, Gzmb*), activation (e.g., *Icos, Tnfrsf4* (OX40), *Tnfrsf9* (4-1BB), *Tnfrsf18* (GITR)), and inhibitory signaling (e.g., *Tigit*, *Lag3, Havcr2* (Tim-3), *Pdcd1* (PD-1)) (**Figure 3a-b**). These genes upregulation underscores their distinct roles in immune evasion and sustaining a suppressive TME, compared to healthy splenic Treg cells. Markers such as GITR and CTLA-4 on TI-Treg cells in liver metastases suppress anti-tumor T cell responses (**Figure 3a**) ^37,41,42^. Subtle differences between liver and peritoneal TI-Treg cells, compared to similarities in primary liver tumor transplants and metastatic liver tumors (**Figure 3c-d**), indicate that TI-Treg cells adapt to the specific tumor environments and highlighting how tumor cells and host tissues shape their phenotypes ^37^. The Wnt signaling pathway is specifically upregulated in peritoneal TI-Treg cells compared liver TI-Treg cells, potentially contributing their functional properties. Sustained Wnt signaling in Treg cells drives epigenetic reprogramming, impairing their immunosuppressive function and promoting β-catenin-mediated transcription of pro-inflammatory genes, linked to CRC progression from inflammatory bowel disease ^43–45^. Moreover, in CRC, enhanced Wnt signaling mediates immune exclusion by increasing Treg cell infiltration and survival while inhibiting CD8^+^ T cell infiltration. Upregulation of Wnt pathway genes has also been linked to Treg cells-enriched tumors ^46^. These findings may explain the accumulation and activation of the Wnt pathway in TI-Treg cells within peritoneal metastases, potentially shaping their functional properties and facilitating CRC progression.

Recent evidence challenges the notion that effector Treg cells adapt to specific tissues and develop unique functions. Instead, tissue Treg cells share a common phenotype and TCR repertoire, allowing migration between tissues ^14^. This complicates the therapeutic aim of selectively targeting TI-Treg cells specifically without altering systemic Treg cell populations. The spleen serves as a key reservoir for circulating and tumor-infiltrating immune cells ^47^. Tumors disrupt splenic immune balance through host and tumor-derived factors that upregulate immune-related functions such as increased myeloid-derived suppressor cell activity ^48–50^. Studies have shown that Treg cell accumulation in tumor-bearing mice increases in draining lymph nodes and spleen, primarily through the conversion of naive CD4^+^ T cells, without thymic involvement or Treg cell proliferation ^51,52^. Furthermore, Valzasina *et al*. demonstrated that Treg cells from the lymph nodes and spleen of tumor-bearing mice resemble natural Treg cells from healthy mice but exhibit an effector-memory phenotype, characterized by reduced CD62L (*Sell*) and increased CD103 (*Itgae*) expression ^52^. These findings align with our data (not shown) and suggest distinct Treg cell subsets circulating between tissues and lymphoid organs ^52^. This accumulation correlates with impaired anti-tumor immunity, allowing tumor evasion of immune responses ^51^. Despite these findings, there was previously a lack of direct comparisons between splenic Treg cells from tumor-bearing mice and TI-Treg cells remains. Deconvolution analysis revealed that tumor-bearing splenic Treg cells form a distinct population, bearing 15% of unique profiles compared to healthy splenic Treg cells and TI-Treg cells from liver tumors (**Figure 6b**). The 85% similarity likely reflects their shared cell type and tissue origin. Our findings support the concept that patients exhibit a systemic response to tumor deveopment, with tumor-bearing splenic Treg cells expressing tumor-associated markers (**Figure 5d**). These results support the importance of evaluating tumor-bearing splenic Treg cells as a reference, to aid in the identification of markers for specifically targeting TI-Treg cells in therapies to minimizing off-target effects.

In summary, this study advances our understanding of TI-Treg cell phenotype and transcriptomic characteristics in CRC metastatic sites. Using the Foxp3EGFP mouse model, we identified differences in TI-Treg cells between liver and peritoneal tumors, with enhanced Wnt signaling prominent in peritoneal tumors. Tumor-bearing splenic Treg cells emerge as a distinct population with unique features. Using these cells as reference for characterizing TI-Treg cells may help identify key transcriptional pathways and signaling mechanisms to target Treg-mediated immune suppression in CRC. This model and approach offers strategies for biomarker discovery and therapies that selectively target tumor-associated Treg cells while preserving systemic immune regulation.

## Acknowledgments

The authors acknowledge Liza Wijler for supplying the mouse tumor organoids and teaching the implantation technique. We also extend our gratitude to the members of the Coffer, Prekovic, Kranenburg and Westendorp research groups for their valuable discussions. We appreciate the support from GDL Utrecht employees for assistance with the mice. Additionally, we thank the Hubrecht Institute FACS facility, Single Cell Discoveries B.V., and the R2 platform. All illustrations were generated using BioRender.com. This work was supported by a Worldwide Cancer Research grant (Reference: 19-0371). The funding agencies played no role in the design, reviewing, or writing of the manuscript.

## Author contributions

S.A.R. designed and conducted experiments, analyzed data, prepared figures, and wrote the manuscript. S.A.R. and A.V. conducted the *in vivo* study from implantation to harvest. C.L.F. performed tissue preparation and IHC staining. Y.K. assisted with deconvolution analysis and participated in writing. B.J.V. assisted with IHC scans analysis. B.G. compiled datasets for survival analysis. E.M. and O.K. supervised the research, contributed to experimental design, and manuscript conceptualization. S.P. assisted with RNA sequencing analysis, manuscript conceptualization, and writing. P.J.C. supervised the research, contributed to experimental design and manuscript conceptualization, and participated in writing.

## Declaration of interest

The authors declare no conflicts of interest.

## Material and methods

### Ethical guidelines

This study was conducted in accordance with institutional guidelines for the care and use of laboratory animals. All animal procedures related to the purpose of the research were approved by the Animal Welfare Body under the Ethical license of University Utrecht, Medical Center Utrecht, The Netherlands, as filed by the relevant national authority, ensuring full compliance with the European Directive 2010/63/EU for the use of animals for scientific purposes.

### Mice

Transgenic B6-Foxp3^EGFP^ male mice (B6.Cg-*Foxp3*^*tm*2*(EGFP)Tch*^/J, The Jackson Laboratory, stock no. 006772) were used in this study. These mice express enhanced green fluorescent protein (eGFP) under the control of the Foxp3 promoter, which is specific to Treg cells. This allows eGFP to serve as a surrogate marker for Treg cells (Foxp3 eGFP), enabling rapid quantification of Treg cell numbers via flow cytometry analysis ^13^. The mice were bred at the Central Laboratory Animal Research Facility (Utrecht) and were housed in groups within open cages, provided with contact bedding, plastic enrichment shelters, and nesting materials. They were fed AIN-93M pellets (Ssniff Spezialdiäten GmbH, Soest, Germany) ad libitum, had unrestricted access to water, and were maintained on a 12-hour light/12-hour dark cycle at a temperature of 20–24°C and a relative humidity of 45–60%. The mice were used for experiments at 10–14 weeks of age.

### CRC tumor-organoid culture

The murine CRC tumor-organoid line was derived from spontaneous colon tumors originated in a transgenic mouse model with conditional activation of the Notch1 receptor and deletion of p53 in the digestive epithelium (NICD/p53^−/-^) ^27,53^. Exome sequencing revealed mutations in either the *Ctnnb1* or *Apc* genes, indicating classical Wnt pathway activation ^53^. Murine CRC tumor-organoids were passaged once a week and medium was refreshed every three days. To passage CRC tumor-organoids, BME (R&D systems) was dispersed by pipetting and washed with pre-cold PBS. CRC tumor-organoids were resuspended thoroughly with pre-warmed TrypLE-Express (Gibco), incubated for 5 minutes at 37°C and mechanically sheared to obtain a single cell suspension. Cells were then washed and resuspended in 50% BME and 50% medium to the right ratio for passaging (1:25). Cells were plated as droplets in pre-warmed culture plate and incubated upside down for approximately 30 minutes at 37°C and 5% CO_2_. After matrix solidification, cells were overlaid with medium. The medium for murine CRC tumor-organoids culture consisted of Advanced Dulbecco’s Modified Eagle Medium (DMEM)/F12 medium (Gibco) supplemented with 10 mM N’-2-Hydroxyethylpiperazine-N’-2 ethanesulphonic acid (HEPES, Lonza), 2 mM Glutamax (Gibco), 50 U/ml penicillin-streptomycin (Gibco), 100 ng/ml Noggin conditioned medium (produced by lentiviral transfection), 1 mM N-acetylcysteine (Sigma) and 2% B27 serum free supplement (Gibco) and 10 nM murine recombinant fibroblast growing factor (PeproTech).

To prepare CRC tumor organoids for orthotopic liver implantation, the organoids were disassociated using TrypLE three days prior to implantation. On the day before the procedure, 100 000 of two-day-old organoids were embedded in 10 µL of 80% Rat Tail Type I Collagen (Corning) droplets mixed with 20% neutralization buffer (Alpha MEM powder (Gibco), 1 M HEPES buffer pH 7.5 (Lonza), and NaHCO3 (Sigma)).

### Microsurgical implantation of tumor organoids into the liver

Orthotopic liver implantation was performed as following the method described by L. Wijler et al. ^27^. Briefly, mice were anesthetized (isoflurane 4% for induction and 2% for maintenance, with 1.6 L/min oxygen) and received subcutaneous buprenorphine (0.1 mg/kg) for analgesia. Under light microscopy and using microsurgical equipment, a small incision was made in the skin and peritoneum to expose the liver, where a 2 mm incision was then created. To control bleeding, a cotton tip was applied while a droplet of collagen containing overnight-recovered CRC tumor organoids was air-dried. Once prepared, the collagen droplet was placed into the incision, which was then sealed with Seprafilm (Genzyme). The peritoneum and skin were sutured, and mice were monitored during recovery and for two days post-surgery to ensure proper wound closure. Following surgery, body weight and tumor progression were monitored weekly through clinical signs and bioluminescent imaging.

### Procedure for euthanization and organ-tumor harvesting

Mice exhibiting signs of tumor growth and metastasis, confirmed through clinical observation and bioluminescence imaging, were euthanized under anesthesia by cervical dislocation. Relevant organs were then harvested post-mortem for further analysis.

### Bioluminescence imaging

Bioluminescence imaging (BLI) was used to monitor *in vivo* tumor progression and metastases formation. Mice were anesthetized and intraperitoneally injected with 100 µL of D-luciferin in PBS, followed by 5-minute imaging (1-second exposure per image) using the PhotonIMAGERTM RT system (Biospace Lab, Paris, France). Following euthanasia, each organ was imaged *ex vivo* to determine the presence and distribution of tumors.

### Immunohistochemistry

Tumor-bearing tissues were fixed in formalin and embedded in paraffin (FFPE). Paraffin blocks were sectioned at 4 µm and mounted onto slides. Before staining, the tissue sections were deparaffinized by immersing in xylene for 10 minutes, followed by rehydration through a graded ethanol series (100% to distilled water). Endogenous peroxidase activity was blocked for 15 minutes, and antigen retrieval was performed by boiling the slides in citrate buffer for 20 minutes. After cooling to room temperature, the slides were washed in PBST for 5 minutes. The sections were stained with a primary anti-mouse CD45 (BD Bioscience) antibody for 2 hours at room temperature, shielded from light. After a 5-minute wash in 0.1% PBS-Tween, the slides were incubated with the secondary HRP Goat Anti-Rat antibody (ImmPRESS, Vector Laboratories Inc) for 1 hour at room temperature, followed by another 5-minute wash in PBS-Tween. Subsequently, the slides were incubated with 3,3′-diaminobenzidine tetrahydrochloride (DAB) for 10 minutes and counterstained with Mayer’s Hematoxylin for 5 seconds. The slides were then air-dried and mounted with ClearVue Mountant XYL (Epredia). Slides were digitally scanned at x40 magnification using the NanoZoomerXR (Hamamatsu) with a resolution of 0.25 µm/pixel, and the resulting scans were analyzed using QuPath software (v0.5.0). Tumour and healthy areas were visually identified, and positive cells were quantified using the positive cell detection tool. The analysis was conducted with the Optical Density Sum for Nuclei detection tool with a pixel size of 0.5 µm, a sigma of 1.3, and a minimum size of 5µm and maximum size of 200 µm. Positive cells were determined on a single threshold using nucleus DAB OD mean to set the cut-off value, allowing for the assessment of DAB-positive cells within both the hotspot areas in a staining specific manner.

### Preparation of single-cell suspensions

The spleen, liver and lungs were harvested individually and smashed against the mesh (70 µm) of a sterile cell strainer in a cell-culture dish containing ice-cold MACs buffer (2% heat-inactivated FBS (Gemini Bio-Products), 2mM EDTA (Sigma-Aldrich) in PBS (Lonza)).

Tumor tissues (0.5-1 g) were collected in cold PBS and transferred to a petri dish with 1 mL of cold RPMI-1640. The tissue was minced into 2-4 mm pieces using a blade and then transferred to a gentleMACS C-Tubes (Miltenyi Biotec). Tissue dissociation was carried out using the murine tumor dissociation kit (Miltenyi Biotec) following the manufacturer’s protocol. The tube was processed in the gentleMACS Octo Dissociator with Heaters (Miltenyi Biotec) using program “37C_h_TDK_1”, which is recommended for soft to medium murine tissues. For tougher tumors with larger pieces remaining, the program “m_imptumor_01” was run for an additional minute. The dissociated tissue was centrifuged at 400g for 1 minute and filtered through a 70 µm strainer. The cell pellet was resuspended in 5 mL RPMI-1640, treated with 200 U/mL DNase I for 5 minutes at room temperature, and then washed.

### Flow cytometry analysis

Single-cell suspensions were initially stained with a live/dead dye, either DAPI (5 minutes) or Zombie NIR (15 minutes), in PBS both at room temperature and shielded from light. This was followed by staining with fluorochrome-labeled antibodies: anti-CD45.2-Pacific blue (BioLegend), anti-CD3-FITC (BioLegend), anti-CD4-APC (Immunotools), anti-CD8-BV570 (BioLegend), anti-CD19(BCR)-BV711 (BioLegend), anti-NK1.1-APC (BioLegend) and anti-CD11B-PE (BioLegend) in MACs buffer for 20 minutes at room temperature, shielded from light. Flow cytometry data were acquired using a BD LSRFortessa Cell Analyzer (BD Biosciences) with FACSDiva (BD Biosciences) software. The data were analyzed with FlowJo (v.10.10.0, Treestar).

### RNA-sequencing and analysis

For RNA sequencing experiments, single-cell suspensions were initially stained with DAPI, as described before, to identify live and dead cells. Subsequently, the cells were stained with anti-CD3-PE (BioLegend) and anti-CD4-APC eFluor780 (Invitrogen) in MACs buffer for 20 minutes at room temperature, shielded from light. Viable cells were then sorted based on CD4 and Foxp3 eGFP expression using a BD FACSAria II and collected into 1.5 mL Eppendorf tubes. Sorted cells were lysed in 100 µl TRIzol (Invitrogen) and stored at -80°C until shipment on dry ice to Single Cell Discoveries B.V. (Utrecht, The Netherlands). RNA extraction was performed using the standard protocol, followed by bulk-cell RNA sequencing using a modified CELSeq protocol ^54,55^. Briefly, RNA samples were barcoded with CEL-seq primers during reverse transcription ^54,55^.

Barcoded cDNA was pooled and linearly amplified in an *in vitro* transcription. Following amplification, RNA was fragmented, and quality was assessed using an Agilent bioanalyzer before library preparation. Sequencing libraries were generated through another round of reverse transcription and PCR amplification to add Illumina Truseq small RNA primers. The quality of the cDNA library was re-evaluated before paired-end sequencing on an Illumina NextSeq 500 System, targeting a depth of 10 million reads per sample. Analysis began with the identification of the Illumina library index and CEL-Seq sample barcode. Sequencing reads were aligned to the mm10 mouse transcriptome (RefSeq: NM_016701.3) using the Burrows-Wheeler Alignment (BWA) tool ^56^. Mapping and count table generation were performed with the MapAndGo script (https://github.com/anna-alemany/transcriptomics/tree/master/mapandgo). Counts were normalized for sequencing depth and RNA composition using DESeq2’s median of ratios method ^57^.

The RNA sequencing data were further analyzed using DESeq2 within R. This included Principal Component Analysis (PCA) with log2 z-score transformation, identification of differentially expressed genes (DEGs), using a significance threshold of p ≤ 0.05, and heatmap generation for gene sets based on mean z-scores. Multiple testing corrections were applied using the false discovery rate (FDR) method with a p-value cutoff of ≤ 0.05 (ANOVA or t-test). Pathway analysis of DEGs was performed in R using appropriate packages (GSVA and clusterProfiler) focusing on gene ontology annotations for biological processes ^58,59^. For Treg-associated pathways, the following datasets were selected: MM7810, MM4647, MM4605, MM1509, MM1411, MM1416, MM1512, MM13149, MM3896, MM732 and MM15404.

To generate 1000 random mixtures, we first sampled 1000 sum-to-one mixing coefficients at random. These samples were projected into PCA space using the principal components trained without incorporating the synthetic mixtures. For deconvolution analysis, we held out two healthy splenic Treg cell samples and five TI-Treg cells from primary liver tumor transplants for evaluation. The remaining samples were used to generate 1000 random mixtures. Subsequently, the held-out samples were signature into a non-negative least square (NNLS) deconvolution analysis to fit both the random mixtures and tumor-bearing splenic Treg cell samples. The deconvolution performance was assessed by calculating the coefficient of determination (R²) between the true profiles and the NNLS-reconstructed profiles. All analyses were conducted in R (version 4.4.2).

### Clinical cohort of colon cancer

Colon cancer cohorts were identified using publicly available databases, specifically the National Center for Biotechnology Information (NCBI) Gene Expression Omnibus (GEO, https://www.ncbi.nlm.nih.gov/geo/) and the Genomic Data Commons (GDC) Data Portal (https://portal.gdc.cancer.gov/). Only datasets with available overall survival and transcriptome-level data were considered, and an initial inclusion criterion was established to focus on cohorts comprising at least thirty patients. To create a unified dataset, duplicate gene arrays were excluded, leading to some cohorts having fewer patients in the final database. To ensure data uniformity and address potential inconsistencies due to variations in sensitivity, specificity, and dynamic range among different gene expression detection platforms, we restricted our selection to tumor samples processed using specific *in-situ* oligonucleotide microarray platforms. These included GPL96 (Affymetrix Human Genome U133A Array), GPL571 (GeneChip Human Genome U133A 2.0 Array), and GPL570 (Affymetrix Human Genome U133 Plus 2.0 Array). These platforms were chosen for their use of identical probe sequences to measure the expression of individual genes, enhancing comparability across datasets. Of note because the GPL96 arrays have less probe set, not each gene is available for each patient sample.

The gene expression data from the selected arrays underwent a two-step normalization procedure. First, MAS5 (Microarray Suite 5.0) normalization was applied, followed by scaling normalization to adjust the mean expression value to 1,000 within each array. To minimize platform-specific discrepancies, only probes present on the GPL96 platform were included in the analysis. This approach was particularly relevant for the GPL570 arrays, which contain a larger number of additional probes. For the selection of the most representative probe set per gene, the JetSet algorithm was utilized to identify the optimal probe based on reliability and specificity ^60^. Finally, matching to between the gene lists and the integrated database was made using the gene symbols approved by the HUGO Human Gene Nomenclature Committee (www.genenames.org).

### Statistical analysis

For FACS data, analysis was performed using GraphPad Prism 10, with results presented as mean values with corresponding standard deviations (SD). Statistical significance was determined using one-way ANOVA with Tukey’s multiple comparison test or unpaired Student’s *t*-test, as appropriate. Immunohistochemistry data were analyzed using unpaired one-way ANOVA with Tukey’s multiple comparison test. Activity scores were compared using non-parametric test (two-tailed). Significance levels are denoted by asterisks: *P* ≤ 0.05 (*), *P* ≤ 0.01 (**), or *P* ≤ 0.001 (***) or *P* < 0.0001 (****).

### Data availability statement

All data supporting the findings of this can be obtained from the lead contact upon request. The RNA sequencing data analyzed in this manuscript have been submitted to GEO and are available to the public as of the publication date. Accession number: GSE275756.

## Notes

### Competing Interest Statement

The authors have declared no competing interest.

