## Supplementary figures and images for "Site-dependent Treg cell transcriptional reprograming in a metastatic colorectal cancer model holds prognostic significance"

### supplementary figures

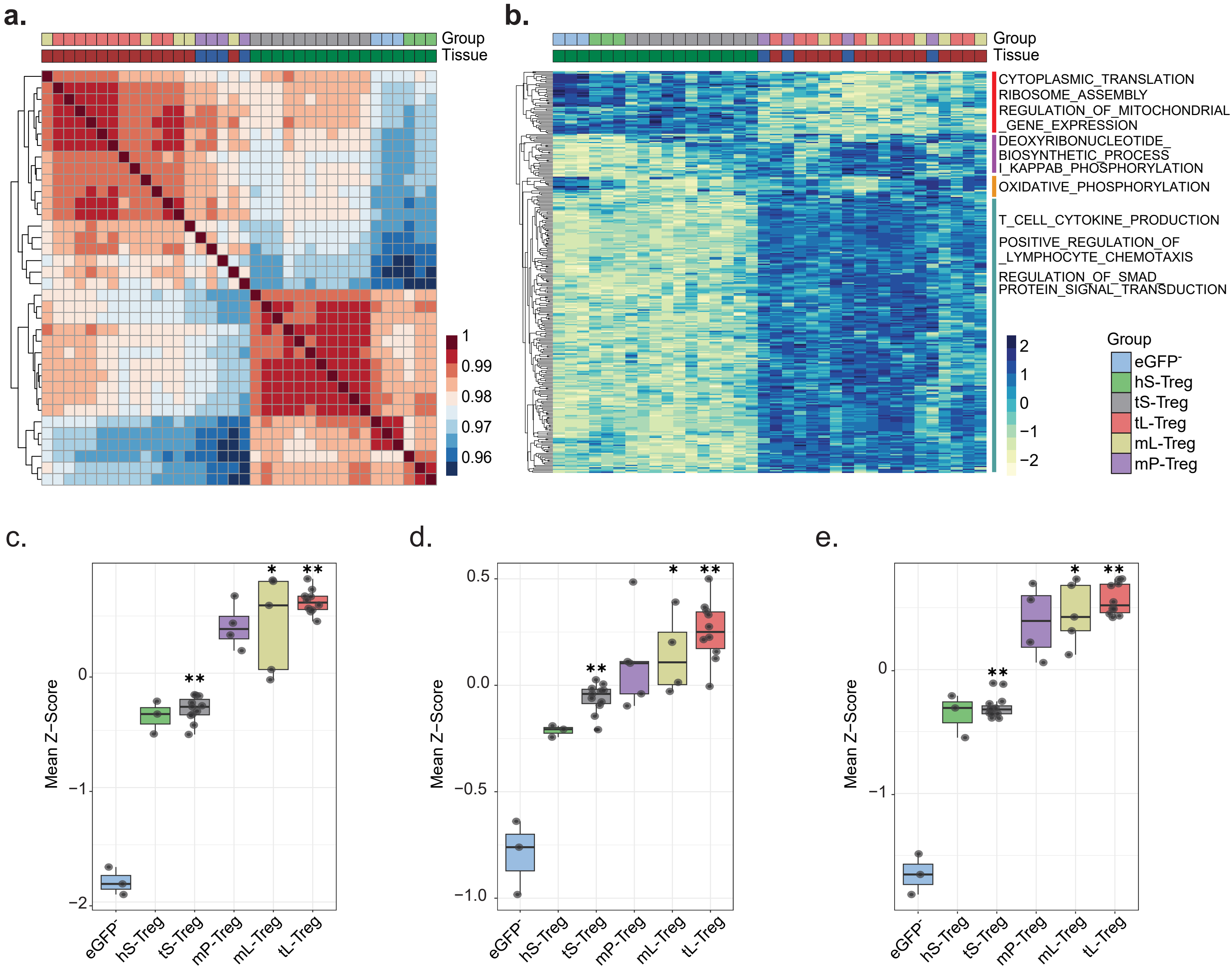


Supplemental Figure 2

Aristin Revilla S. *et al.*

### supplementary figures legends

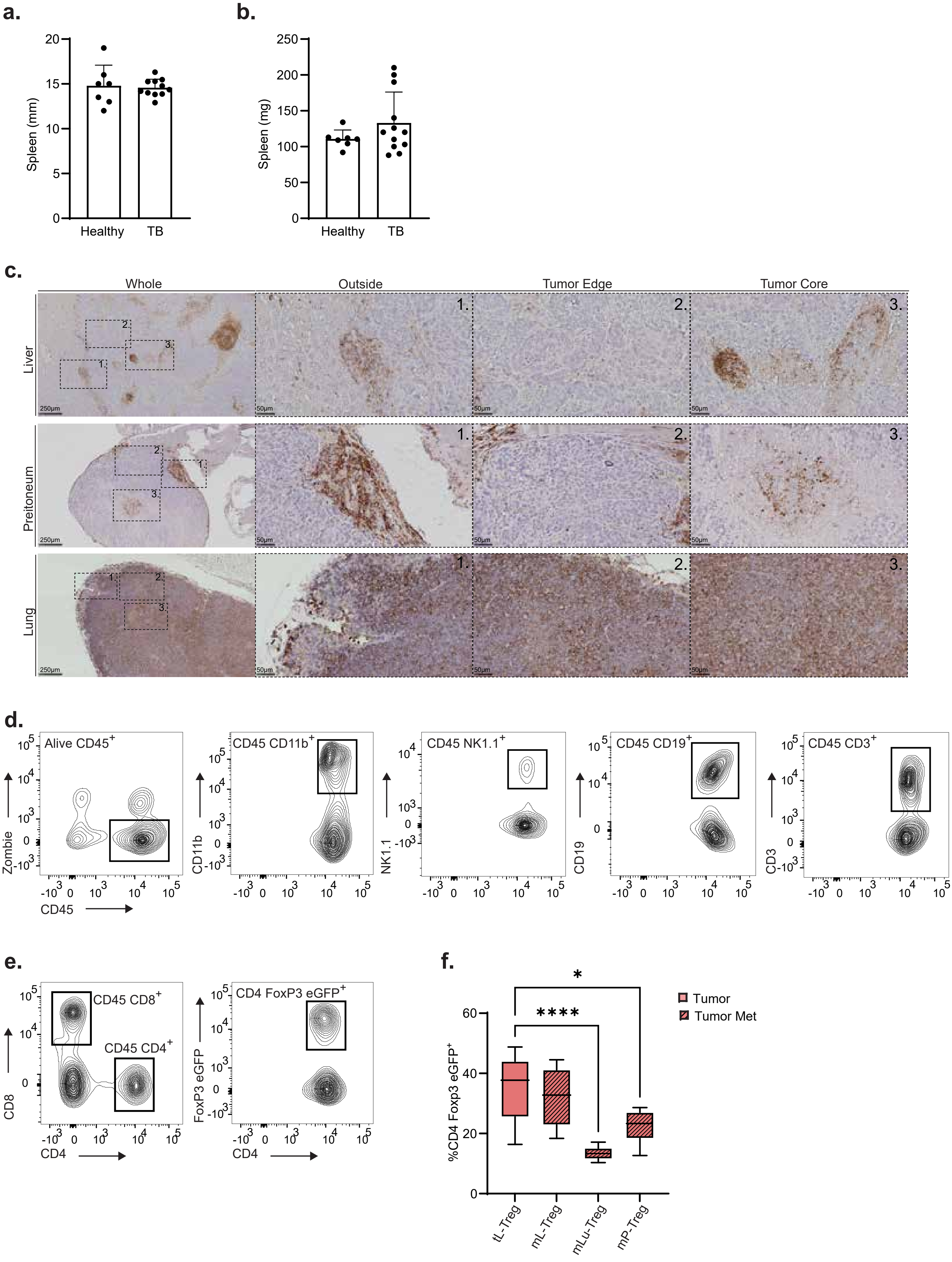
